# Equivalent high-resolution identification of neuronal cell types with single-nucleus and single-cell RNA-sequencing

**DOI:** 10.1101/239749

**Authors:** Trygve E. Bakken, Rebecca D. Hodge, Jeremy M. Miller, Zizhen Yao, Thuc N. Nguyen, Brian Aevermann, Eliza Barkan, Darren Bertagnolli, Tamara Casper, Nick Dee, Emma Garren, Jeff Goldy, Lucas T. Gray, Matthew Kroll, Roger S. Lasken, Kanan Lathia, Sheana Parry, Christine Rimorin, Richard H. Scheuermann, Nicholas J. Schork, Soraya I. Shehata, Michael Tieu, John W. Phillips, Amy Bernard, Kimberly A. Smith, Hongkui Zeng, Ed S. Lein, Bosiljka Tasic

## Abstract

Transcriptional profiling of complex tissues by RNA-sequencing of single nuclei presents some advantages over whole cell analysis. It enables unbiased cellular coverage, lack of cell isolation-based transcriptional effects, and application to archived frozen specimens. Using a well-matched pair of single-nucleus RNA-seq (snRNA-seq) and single-cell RNA-seq (scRNA-seq) SMART-Seq v4 datasets from mouse visual cortex, we demonstrate that similarly high-resolution clustering of closely related neuronal types can be achieved with both methods if intronic sequences are included in nuclear RNA-seq analysis. More transcripts are detected in individual whole cells (∼11,000 genes) than nuclei (∼7,000 genes), but the majority of genes have similar detection across cells and nuclei. We estimate that the nuclear proportion of total cellular mRNA varies from 20% to over 50% for large and small pyramidal neurons, respectively. Together, these results illustrate the high information content of nuclear RNA for characterization of cellular diversity in brain tissues.

## Introduction

Understanding neural circuits requires characterization of their cellular components. Cell types in mammalian brain have been defined based on shared morphological, electrophysiological and, more recently, molecular properties (Poulin et al. 2016; Zeng and Sanes 2017; Bernard, Sorensen, and Lein 2009). scRNA-seq has emerged as a high-throughput method for quantification of the majority of transcripts in thousands of cells. scRNA-seq data have revealed diverse cell types in many mouse brain regions, including neocortex (Tasic et al. 2016; Tasic et al. 2017; Zeisel et al. 2015), hypothalamus (Campbell et al. 2017), and retina (Shekhar et al. 2016; Macosko et al. 2015).

However, scRNA-seq profiling does not provide an unbiased survey of neural cell types. Some cell types are more vulnerable to the tissue dissociation process and are underrepresented in the final data set. For example, in mouse neocortex, fast-spiking parvalbumin-positive interneurons and deep-projecting glutamatergic neurons in layer 5b are observed in lower proportions than expected and need to be selectively enriched using Cre-driver lines (Tasic et al. 2017) for sufficient sampling. In adult human neocortex, neurons largely do not survive dissociation thereby causing over-representation of non-neuronal cells in single cell suspensions (Darmanis et al. 2015). In contrast to whole cells, nuclei are more resistant to mechanical assaults and can be isolated from frozen tissue (Krishnaswami et al. 2016; Lacar et al. 2016). Single nuclei have been shown to provide sufficient gene expression information to define relatively broad cell classes in adult human brain (Lake et al. 2016; Lake et al. 2017) and mouse hippocampus (Habib et al. 2016).

Previous studies have not addressed if the nucleus contains sufficient diversity and number of transcripts to enable discrimination of closely related cell types at a resolution comparable to whole cells. A recent study compared clustering results for single nuclei and whole cells isolated from mouse somatosensory cortex (Lake et al. 2017), but it only showed similar ability to distinguish two very different cell classes: superficial- and deep-layer excitatory neurons.

In this study, we compared 463 matched nuclei and whole cells from layer 5 of mouse primary visual cortex (VISp) to investigate differences in single nucleus and single cell transcriptomes. We selected this brain region because it contains a known variety of distinguishable yet highly similar cell types that would reveal the cell-type detection limit of RNA-seq data obtained from single cells or nuclei (Tasic et al. 2016). We used the same primary cell source and processed cells and nuclei with the same transcriptomic profiling method to directly compare the resolution limit of cell type detection from well-matched sets of single cells and nuclei. Furthermore, we compared the nuclear fraction of gene transcripts among cell types and identified functional classes of transcripts that are enriched in the cytoplasm and nucleus.

## Results

### RNA-seq profiling of single nuclei and single cells

We isolated 487 NeuN-positive single nuclei from layer 5 of mouse VISp using fluorescence activated cell sorting (FACS). Anti-NeuN staining was performed to enrich for neurons. In parallel, we isolated 12,866 tdT-positive single cells by FACS from all layers of mouse VISp and a variety of Cre-driver lines, as part of a larger study on cortical cell type diversity (Tasic et al. 2017). For both single nuclei and cells, poly(A)-transcripts were reverse transcribed and amplified with SMART-Seq v4, cDNA was tagmented by Nextera XT, and resulting libraries were sequenced to an average depth of 2.5 million reads (Figure 1A). RNA-seq reads were mapped to the mouse genome using the STAR aligner (Dobin et al. 2013). Gene expression was quantified as the sum of intronic and exonic reads per gene and was normalized as counts per million (CPM) and log2-transformed. For each nucleus and cell, the probabilities of gene detection dropouts were estimated as a function of average expression level based on empirical noise models (Kharchenko, Silberstein, and Scadden 2014).

**Figure 1:**
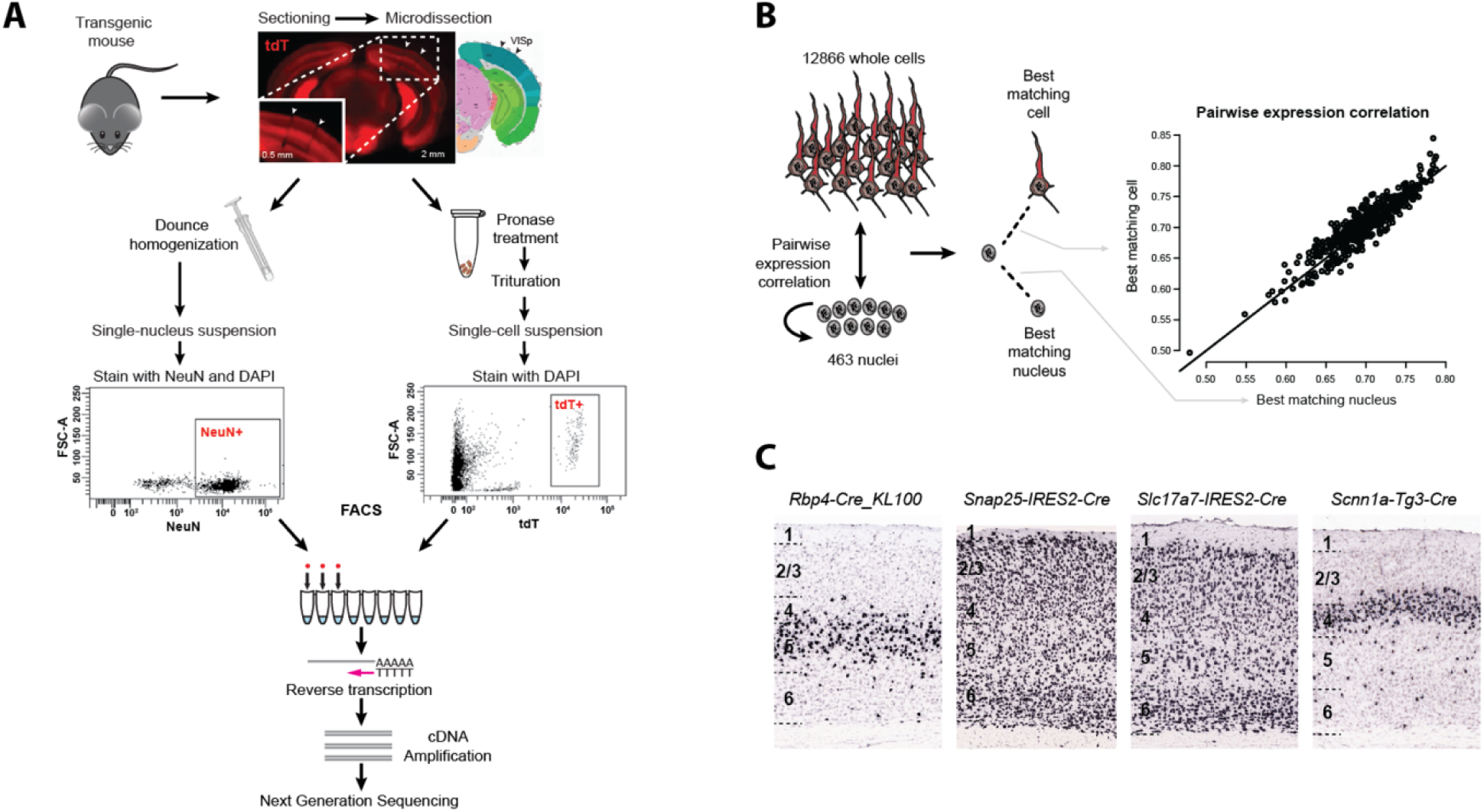
Identification of an expression-matched set of single nuclei and whole cells from mouse primary visual cortex (VISp). **(A)** Whole brains were dissected from transgenic mice, coronal slices were sectioned, and individual layers of VISp were microdissected. Nuclei were dissociated from layer 5, stained with DAPI and against the neuronal marker NeuN. Single NeuN-positive nuclei were isolated by fluorescence-activated cell sorting (FACS). In parallel, whole cells were dissociated from all layers, and single td-Tomato reporter-positive cells were isolated. Single nucleus and cell mRNA were reverse transcribed, amplified, and sequenced to measure transcriptome-wide expression levels. **(B)** Left: 463 nuclei from layer 5 and 12,866 whole cells from all layers passed quality control metrics, and the expression correlation was calculated between each nucleus and all other nuclei and cells. Expression similarity can vary based on sample quality, so nuclei were compared to each other to provide a baseline expected similarity. For each nucleus, the best matching nucleus and cell were selected based on maximal correlation. Right: Cells and nuclei displayed comparable expression similarities to all nuclei, with 95% of correlations between 0.63 and 0.78. This suggested that nuclei and cells were well matched. **(C)** Chromogenic RNA *In situ* hybridization (ISH) images of all VISp layers from four mouse Cre-lines from which the best matching cells were most commonly derived. As expected, all Cre-lines label cells in layer 5 and adjacent layers.

463 out of 487 single nuclei (95%) passed quality control metrics, and each nucleus was matched to the most similar nucleus and cell based on the maximum correlated expression of all genes, weighted for gene dropouts. Nuclei had similarly high pairwise correlations to cells as to other nuclei suggesting that cells and nuclei were well matched (Figure 1B). As expected, matched cells were derived almost exclusively from layer 5 and adjacent layers 4 and 6 (Figure S1B), and from Cre-driver lines that labeled cells in layer 5 (Figure 1C and Figure S1A,C). The small minority of matched cells isolated from superficial layers were GABAergic interneurons that have been detected in many layers (Tasic et al. 2017).

### Comparison of nuclear and whole cell transcriptomes

scRNA-seq profiles nuclear and cytoplasmic transcripts, whereas snRNA-seq profiles nuclear transcripts. Therefore, we expect that RNA-seq reads will differ between nuclei and cells. In nuclei, more than 50% of reads that aligned to the mouse genome did not map to known spliced transcripts but to non-exonic regions within gene boundaries. They were therefore annotated as intronic reads (Figure 2A). In contrast, the majority of cells had less than 30% intronic reads with a minority of cells having closer to 50% intronic reads, similar to nuclei. Median gene detection based on exonic reads was lower for nuclei (∼5,000 genes) than for cells (∼9,500). Including both intronic and exonic reads increased gene detection for nuclei (∼7,000) and cells (∼11,000), demonstrating that intronic reads provided additional information not captured by exons. Whole brain control RNA displayed a read mapping distribution similar to cells, which is consistent with dissociated single cells capturing the majority of transcripts in the whole cell.

**Figure 2:**
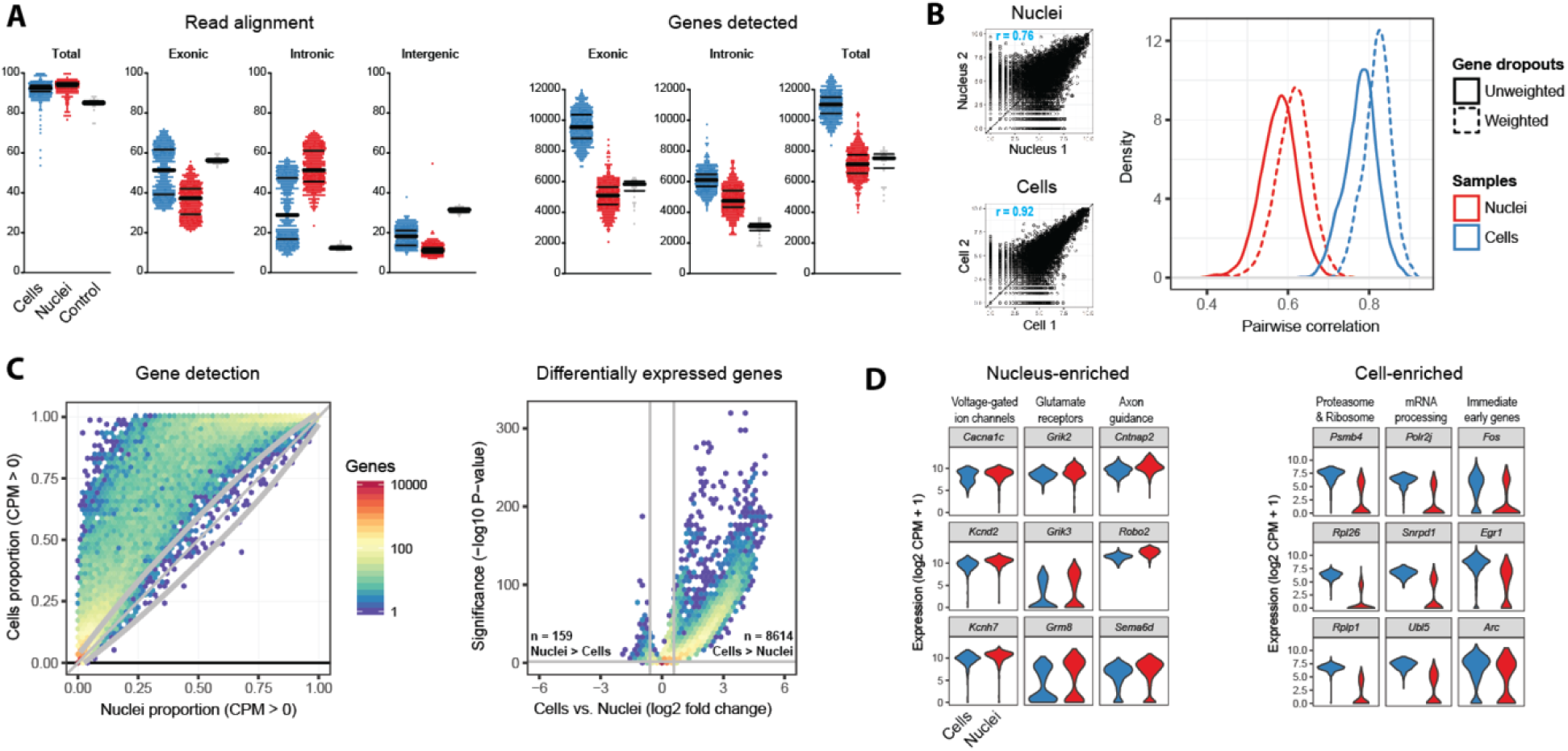
Comparison of nuclear and whole cell transcriptomes. **(A)** Left: Percentage of RNA-seq reads mapping to genomic regions for cells, nuclei, and whole brain control RNA. Bars indicate median and 25^th^ and 75^th^ quantiles. Note that among cells exonic and intronic read alignment is bimodal. Right: Gene detection (counts per million, CPM > 0) based on reads mapping to exons, introns, or both introns and exons. **(B)** Left: The most similar pair of cells have more highly correlated gene expression (r = 0.92) than the most similar pair of nuclei (r = 0.76), due to fewer gene dropouts. Right: Cells have consistently more similar expression to each other than nuclei, even after correcting for gene dropouts based on an expression noise model. **(C)** Left: Binned scatter plot showing all genes are detected (CPM > 0) with equal or greater reliability in cells than nuclei. Grey lines show the variation in detection that is expected by chance (95% confidence interval). Right: Binned scatter plot showing 0.4% of genes are significantly more highly expressed (fold change > 1.5, adjusted P-value < 0.05) in nuclei, and 20.5% of genes are more highly expressed in cells. The log-transformed color scale indicates the number of genes in each bin. **(D)** Nuclear enriched genes are highly enriched for genes involved in neuronal connectivity, synaptic transmission, and intrinsic firing properties. Cell enriched genes are predominantly related to mRNA processing and protein translation and degradation. In addition, immediate early gene expression is increased up to 10-fold in cells, despite comparable isolation protocols for cells and nuclei.

Transcript dropouts likely result from both technical and biological variability, and both effects are more pronounced in nuclei than in cells. When transcript dropouts were adjusted based on empirical noise models, correlations between pairs of nuclei and pairs of cells increased, although cell-cell similarities remained significantly higher (Figure 2B). A majority of expressed genes (21,279; 63%) showed similar detection (<10% difference) in nuclei and cells, whereas 7,217 genes (21%) were detected in at least 25% more cells than nuclei (Figure 2C and Table S1). 8,614 genes have significantly higher expression in cells than nuclei (>1.5 fold expression; FDR < 0.05) and many are involved in house-keeping functions such as mRNA processing and translation (Figure 2D). Genetic markers of neuronal activity, such as immediate early genes *Fos*, *Egr1*, and *Arc* also displayed up to 10-fold increased expression in cells, potentially a byproduct of tissue dissociation (Lacar et al. 2016). 159 genes have significantly higher expression in nuclei (Figure 2D and Table S2), and they appear relevant to neuronal identity as they include connectivity and signaling genes (Figure S2A and Table S4). Based on the sum of intronic and exonic reads, these 159 nucleus-enriched genes are on average more than 10-fold longer than cell-enriched genes (Figure S2B), as recently reported for single nuclei in mouse somatosensory cortex (Lake et al. 2017). When only exonic reads were used to quantify expression in nuclei and cells, a different set of 146 genes were significantly enriched in nuclei (Table S3) and were only slightly longer than cell-enriched genes. These genes were not associated with neuron-specific functions and were significantly enriched for genes that participate in pre-mRNA splicing.

### Intronic reads are required for high-resolution cell type identification from snRNA-seq

Next, we applied an iterative clustering procedure (see Methods and Figure S3) to identify clusters of single nuclei and cells that share gene expression profiles. To assess cluster robustness, we repeated clustering 100 times using 80% random subsets of nuclei and cells and calculated the proportion of clustering runs in which each pair of samples clustered together. Co-clustering matrices were reordered using Ward’s hierarchical clustering and represented as heatmaps with coherent clusters ordered as squares along the diagonal (Figure 3A,Figure B).

**Figure 3:**
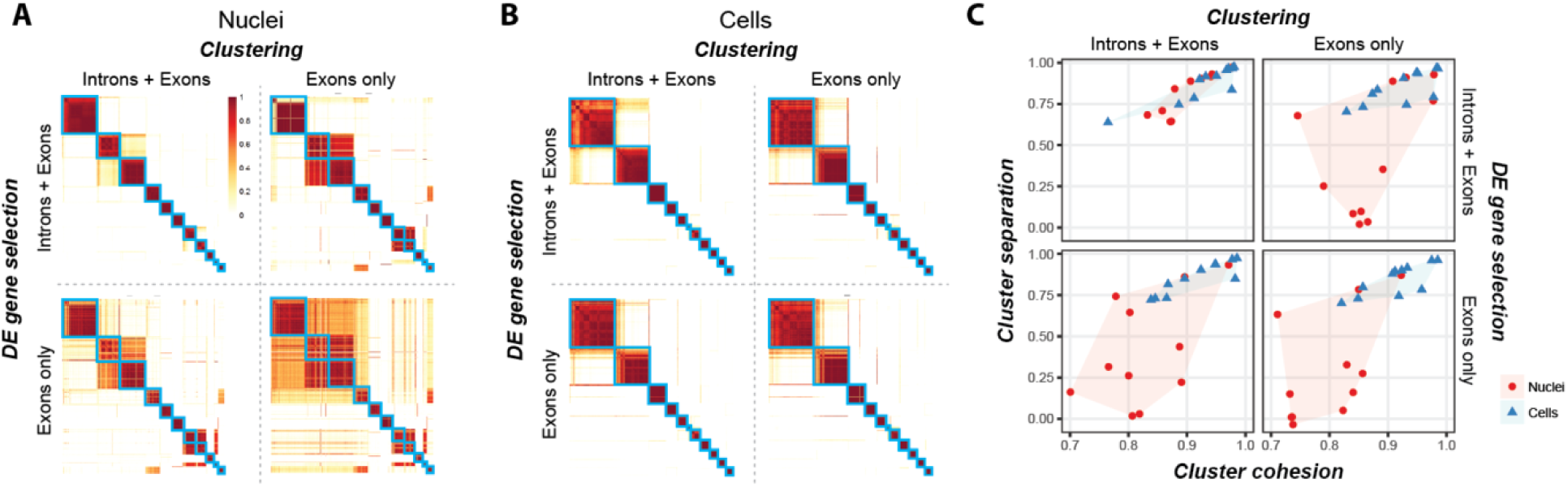
Single nuclei provide comparable clustering resolution to cells with inclusion of intronic reads. **(A)** Co-clustering heatmaps show the proportion of 100 clustering iterations that each pair of nuclei were assigned to the same cluster. Clustering was performed using gene expression quantified with exonic reads or intronic plus exonic reads for two key clustering steps: selecting significantly differentially expressed (DE) genes and calculating pairwise similarities between nuclei. Co-clustering heatmaps were generated for each combination of gene expression values, and blue boxes highlight 11 clusters of nuclei that consistently co-clustered using introns and exons (upper left heatmap) and were overlaid on the remaining heatmaps. The row and column order of nuclei is the same for all heatmaps. **(B)** Co-clustering heatmaps were generated for cells as described for nuclei in **(A)**, and blue boxes highlight 11 clusters of cells. **(C)** Cluster cohesion (average within cluster co-clustering) and separation (difference between within cluster co-clustering and maximum between cluster co-clustering) are plotted for nuclei and cells and all combinations of reads. Including introns in gene expression quantification dramatically increases cohesion and separation of nuclei but not cell clusters.

Clustering includes two steps – differentially expressed (DE) gene selection and distance measurement – that are particularly sensitive to expression quantification. We repeated clustering using intronic and exonic reads or only exonic reads for these steps, and ordered co-clustering matrices to match the results using all reads for both steps. When using introns and exons, we found 11 distinct clusters of nuclei and cells, and clusters had similar cohesion (average within cluster co-clustering) and separation (average co-clustering difference with the closest cluster) (Figure 3C). Including intronic reads for either clustering step increased the number of clusters detected for nuclei but not cells. Therefore, accounting for intronic reads in snRNA-seq was critical to enable high-resolution cluster detection equivalent to that observed with scRNA-seq.

### Equivalent cell types identified with nuclei and cells

We used hierarchical clustering of median gene expression values in each cluster to determine the relationships between clusters. We find that cluster relationships represented as dendrograms are remarkably similar for nuclei and cells (Figure 4A). We compared the 11 clusters identified with single nuclei and cells to reported cell types in mouse VISp (Tasic et al. 2016). Each nucleus and cell cluster could be linked to a reported cell type (Figure S4A) and to each other (Figure 4B) based on correlated expression of marker genes. Many genes contributed to high expression correlations (r > 0.85) for all cluster pairs (Figure S4B). Conserved marker gene expression confirmed that the same 11 cell types were identified with nuclei and cells (Figure 4C). These cell types included nine excitatory neuron types from layers 4-6 and two inhibitory interneuron types. Matched cluster proportions were mostly consistent, except two closely related layer 5a subtypes were under-(L5a Batf3) or over-represented (L5a Hsd11b1) among cells (Figure S4C). This demonstrated that the initial matching of cells to nuclei was relatively unbiased.

**Figure 4:**
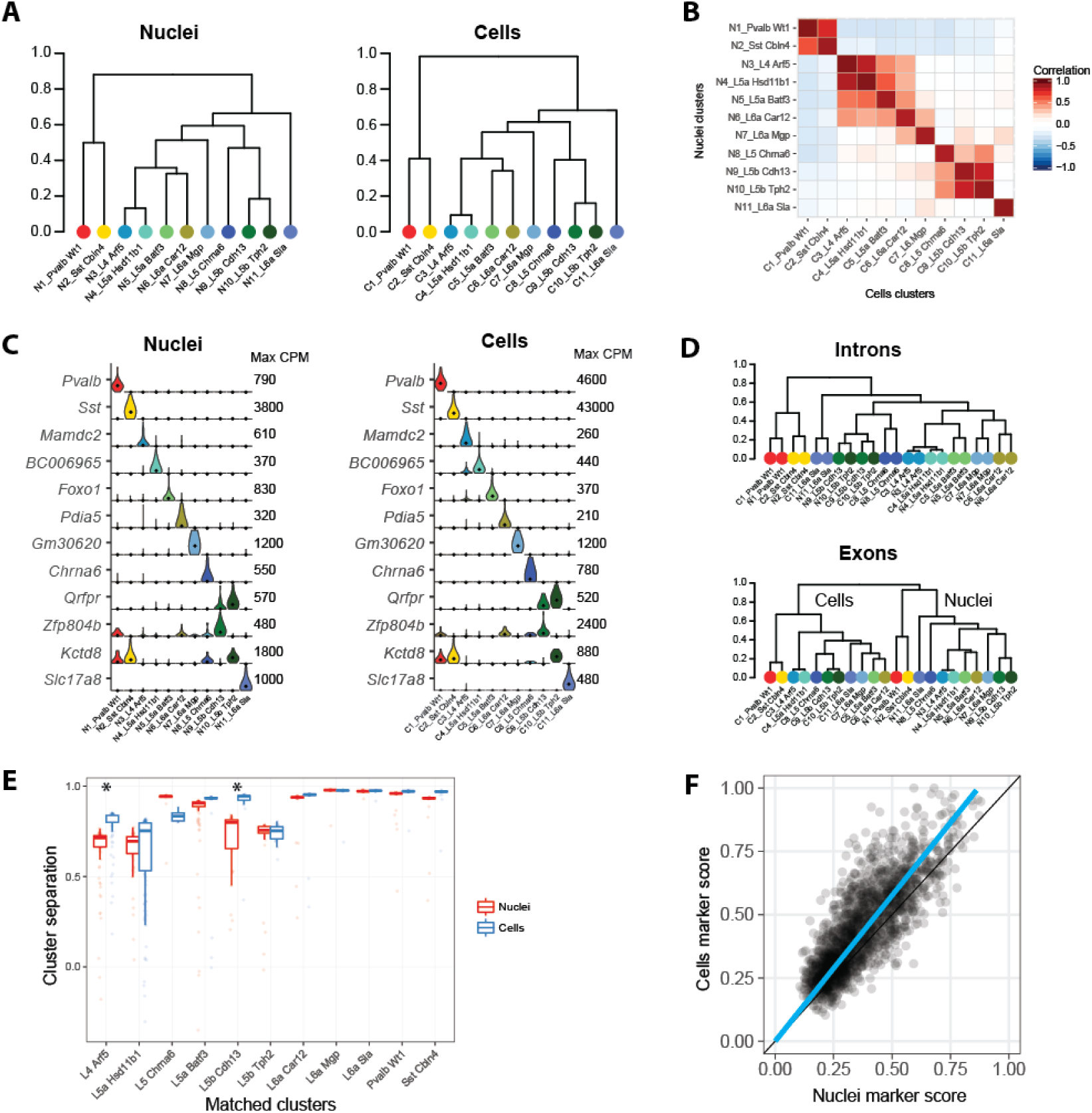
Equivalent neuronal cell types identified with nuclei and cells. **(A)** Cluster dendrograms for nuclei and cells based on hierarchical clustering of average expression of the top 1200 cluster marker genes. 11 clusters are labeled based on dendrogram leaf order and the closest matching mouse VISp cell type described in (Tasic et al. 2016) based on correlated marker gene expression (see Figure S4). **(B)** Pairwise correlations between nuclear and cell clusters using average cluster expression of the top 490 shared marker genes. **(C)** Violin plots of cell type specific marker genes expressed in matching nuclear and cell clusters. Plots are on a linear scale, max CPM indicates the maximum expression of each gene, and black dots indicate median expression. **(D)** Hierarchical clustering of nuclear and cell clusters using the top 1200 marker genes with expression quantified by intronic or exonic reads. Intronic reads group nine matching nuclear and cell clusters together at the leaves, while two closely related deep layer 5 excitatory neuron types group by sample type. In contrast, exonic reads completely segregate clusters by sample type. **(E)** Box plots of cluster separations for all samples in matched nuclear and cell clusters. Clusters are equally well separated for all but two cell types, L4 Arf5 and L5b Cdh13, that are moderately but significantly (Wilcoxon signed rank unpaired tests; Bonferroni corrected P-value < 0.05) more distinct with cells than nuclei. **(F)** Cell type marker genes are consistently detected in nuclei and cells, although marker scores (see Methods) were on average 15% higher for cells.

We hypothesized that most intronic reads were mapped to nuclear transcripts, so quantifying gene expression in cells using only introns would approximate nuclear expression. This was supported by higher correlations of average expression across all nuclei and cells using only intronic reads as compared to only exonic reads (Figure S4D). Thus, a dendrogram based on the median expression (quantified using only intronic reads) of nuclei and cell clusters paired all matching cell types, except for two closely related layer 5b subtypes (Figure 4D). Therefore, intronic reads can help facilitate comparisons between data sets derived from snRNA-seq and scRNA-seq although small expression differences remain. A dendrogram based on exonic reads grouped clusters first by sample type (nuclei and cells) and then by broad cell class (inhibitory and excitatory neurons). Samples grouped by sample type likely due to differences in cytoplasmic transcripts that were profiled in cells but not nuclei. A dendrogram based on intronic reads did not show this grouping because most cytoplasmic transcripts are spliced so were quantified by exonic but not intronic reads.

While we detected the same cell types using nuclei and cells, we expected that gene expression captured with cells included additional information from cytoplasmic transcripts. We compared the separation of matched pairs of clusters based on co-clustering and found that all nuclei and cell clusters were similarly distinct, except using single cells significantly increased the separation of two pairs of similar types: L4 Arf5 from L5a Hsd11b1 and L5b Cdh13 from L5b Tph2 (Figure 4E). Next, we compared how well genes marked cell types by calculating the degree of binary expression. Cell marker scores were, on average, 15% higher than nucleus scores due to fewer expression dropouts in cells (Figure 4F), and this was consistent with mildly improved cluster separation.

### Nuclear content varies among cell types and for different transcripts

We estimated the nuclear proportion of mRNA for each cell type in two ways. Transcripts in the cytoplasm are spliced so intronic reads should be restricted to the nucleus. First, we estimated the nuclear proportion by calculating the ratio of the percentage of intronic reads in cells to the percentage of intronic reads in nuclei (Figure 5A). Second, we estimated nuclear proportions by selecting three genes (*Malat1*, *Meg3*, and *Snhg11*) with the highest expression in nuclei (Figure S4D) and calculating the ratio of the average expression in cells versus nuclei (Figure 5B and Figure S5A). Both methods predicted that L4 Arf5 and L5a Hsd11b1 had a significantly larger proportion of transcripts located in the nucleus compared to other cell types (Figure 5C).

**Figure 5:**
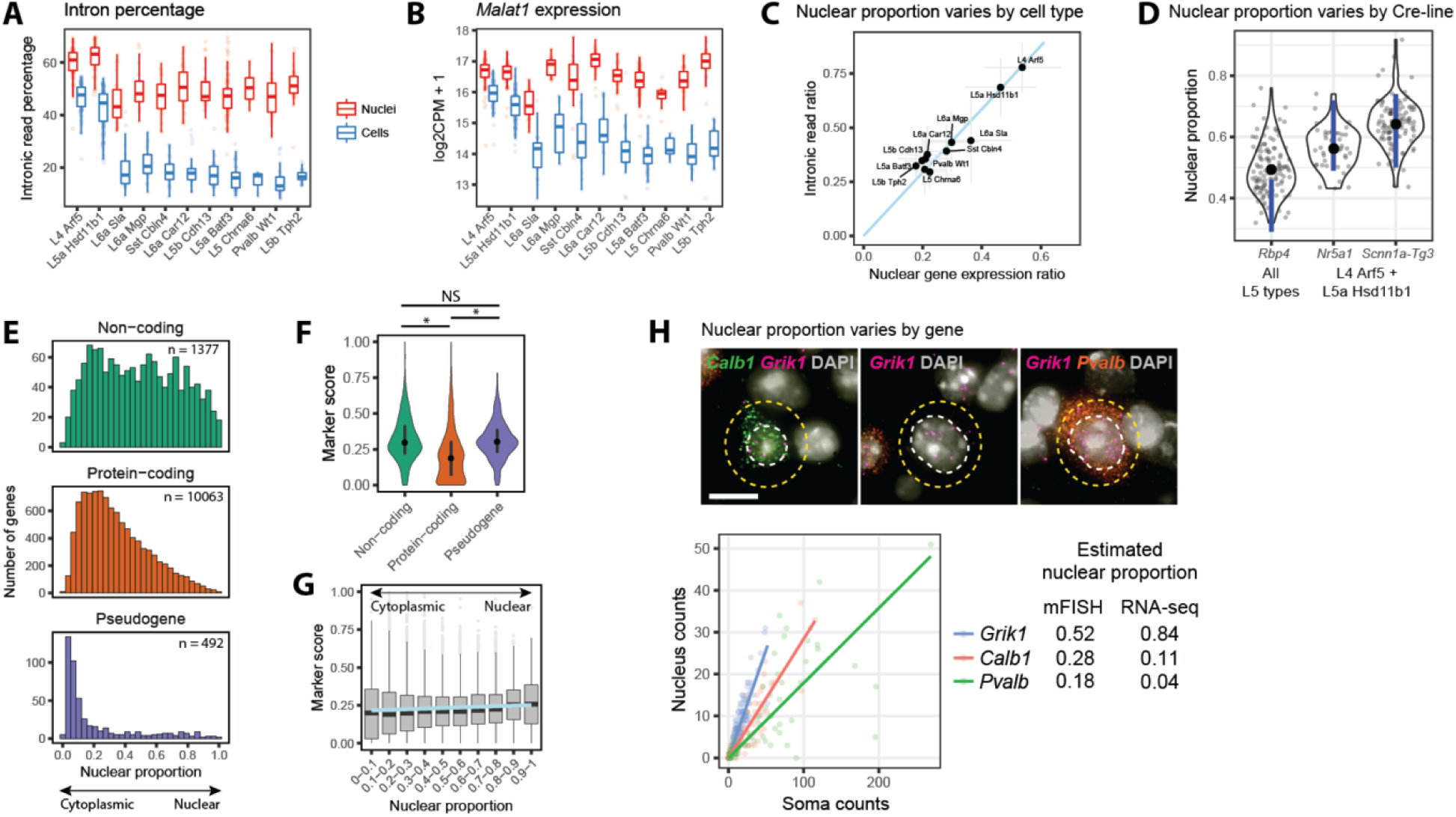
Nuclear transcript content varies among cell types and genes. **(A)** Box plots showing median (bars), 25^th^ and 75^th^ quantiles (boxes), and range (whiskers) of percentages of reads mapping to introns for matched nuclei and cell clusters. **(B)** Box plots of log_2_-transformed expression of the nuclear non-coding RNA, *Malat1*, in matched nuclei and cell clusters. **(C)** The nuclear fraction of transcripts in cell types was estimated with two methods: the ratio of intronic read percentages in cells compared to nuclei; and the average ratio of expression in cells compared to nuclei of three highly expressed genes (*Snhg11, Meg3, and Malat1*) that are localized to the nucleus. The relative ranking of nuclear fractions was consistent (Spearman rank correlation 0.84), although estimates based on the intronic read ratio were consistently 50% higher. **(D)** Estimated nuclear proportion (ratio of nucleus and soma volume) of neurons labeled by three mouse Cre-lines in Layers 4 and 5 (see Supplementary Figure S5D). Single neuron measurements (grey points) were summarized as violin plots, and average nuclear proportions (black points) were compared to the range of estimated proportions (blue lines) based on intronic read ratios and nuclear gene expression. **(E)** Histograms of nuclear fraction estimates for 11,932 genes expressed (CPM > 1) in at least one nuclear or cell cluster and grouped by type of gene. **(F)** Violin plots of marker score distributions with median and inter-quartile intervals. Non-coding genes and pseudogenes are on average better markers of cell types than protein-coding genes. Kruskal-Wallis rank sum test, post hoc Wilcoxon signed rank unpaired tests: *P < 1 x 10^-50^ (Bonferroni-corrected), NS, not significant. **(G)** Box plots of cell type marker scores for genes grouped by estimated nuclear enrichment. Nucleus-enriched genes have significantly higher marker scores (linear regression; P = 2.3 x 10^-8^). **(H)** Validation of the estimated nuclear proportion of transcripts for *Calb1*, *Grik1*, and *Pvalb* using multiplex fluorescent *in situ* hybridization (mFISH). Top: For each gene, transcripts were labeled with fluorescent probes and counted in the nucleus (white) and soma (yellow). Bottom: Probe counts in the nucleus and soma across all cells with linear regression fits to estimate nuclear transcript proportions for each gene. Estimated proportions based on mFISH and RNA-seq data are summarized on the right.

Based on the comparison of scRNA-seq and snRNA-seq data, we estimate that L4 types have high nuclear to cell volume (∼50%), whereas L5 types have lower nuclear to cell volume. To evaluate this finding, we measured nucleus and soma sizes of different cell types *in situ*. These types were labeled by different Cre-transgenes and a Cre-reporter. *Nr5a1*-Cre and *Scnn1a-Tg3*-Cre mice almost exclusively label two cell types (L4 Arf5 and L5a Hsd11b1), whereas *Rbp4*-Cre mice label all layer 5 cell types including L5a Hsd11b1 (Figure S5B and Table S5). We measured the nuclear and cell sizes *in situ*, and calculated the nuclear proportion of each cell as the ratio of nuclear to soma volume (Figure S5C). We found that the average nuclear proportion was significantly lower for layer 5 cells compared to layer 4 cells, as predicted based on RNA-seq data (Figure 5D).

In addition, nuclear proportion estimates based on *in situ* size measurements were systematically higher than predicted for layer 5 but not layer 4 neurons. This could be the result of under-estimating the soma volume based on cross-sectional area measurements of these large non-spherical (pyramidal) neurons. Alternatively, layer 5 neuronal nuclei may have a lower density of nuclear transcripts or there may be cell type-specific biases in our RNA-seq based estimates. We then performed an unbiased survey of nuclear proportions across the full depth of cortex to test whether layer 4 or layer 5 neurons were exceptional compared to neurons in other layers. We found that layer 5 neurons tend to be larger and have proportionally smaller nuclei (Figure S5D) than other cortical neurons, and this feature is also found in rat primary visual cortex (Sigl-Glöckner and Brecht 2017).

Next, we determined the nuclear versus cytoplasmic distribution of transcripts for individual genes. The nuclear proportion of 11,932 transcripts was estimated by the ratio of nuclear to whole cell expression multiplied by the overall nuclear fraction of each cell type and averaged across cell types (Table S6). Different functional classes of genes had strikingly different nuclear proportions (Figure 5E). Many non-coding transcripts were localized in the nucleus, but some were abundantly expressed in the cytoplasm, such as the long non-coding RNA (lncRNA) *Tunar* that is highly enriched in the brain, is conserved across vertebrates, and has been associated with striatal pathology in Huntington’s disease (Lin et al. 2014). Most protein-coding transcripts were expressed in both the nucleus and cytoplasm with a small number restricted to the nucleus, including the Parkinson’s risk gene *Park2*. We found that pseudogenes were almost exclusively cytoplasmic and were highly enriched for house-keeping functions.

We compared our estimates of nuclear enrichment in cortex to mouse liver and pancreas based on data from (Halpern et al. 2015) and found moderately high correlation (r = 0.61) between 4,373 mostly housekeeping genes that were expressed in all three tissues. Moreover, the shape of the distributions of nuclear transcript proportions was highly similar between tissues with slightly higher proportions estimated in this study. These results suggest that the mechanisms regulating the spatial localization of these transcripts – for example, rates of nuclear export and cytoplasmic degradation (Halpern et al. 2015) – are conserved across cell types.

Surprisingly, non-coding genes and pseudogenes are better markers of cell types, on average, than protein-coding genes (Figure 5F). lncRNAs are known to have more specific expression among diverse human cell lines (Djebali et al. 2012), and we show that this is also true for neuronal types in the mouse cortex. Many pseudogene transcripts, most of which are enriched in the cytoplasm, were selectively depleted in the two cell types, L4 Arf5 and L5a Hsd11b1. This is consistent with our previous analysis that showed that neurons of these types have relatively less cytoplasm. We also find that nucleus-enriched transcripts are slightly better cell-type markers than cytoplasm-enriched transcripts, although this is highly variable across genes (Figure 5G).

Finally, we compared our estimates of nuclear localization of transcripts for three genes – *Calb1*, *Grik1*, and *Pvalb* – to relative counts of transcripts in nuclei and cytoplasm using multiplex RNA fluorescence *in situ* hybridization (mFISH). We found that the relative nuclear proportions estimated by scRNA-seq and mFISH were consistent although the absolute levels were quite variable (Figure 5H). Both methods confirmed that *Pvalb* transcripts were mostly excluded from the nucleus, and this explained why 2 out of 35 nuclei in the Pvalb-positive interneuron type (Pvalb Wt1) had no detectable *Pvalb* expression, whereas all cells of this cell type had robust *Pvalb* expression.

## Discussion

Unlike scRNA-seq, snRNA-seq enables transcriptomic profiling of tissues that are refractory to whole cell dissociation and of archived frozen specimens. snRNA-seq is also less susceptible to perturbations in gene expression that occur during cell isolation, such as increased expression of immediate early genes that can obscure transcriptional signatures of neuronal activity (Lacar et al. 2016). However, these advantages come at the cost of profiling less mRNA, and until this study, it was unclear if the nucleus contained sufficient number and diversity of transcripts to distinguish highly related cell types.

To directly address this question, we profiled a well-matched set of 463 nuclei and 463 cells from layer 5 of mouse primary visual cortex and identified 11 matching neuronal types: 2 interneuron types and 9 similar excitatory neuron types. Including intronic reads in gene expression quantification was necessary to achieve high-resolution cell type identification from single nuclei. Intronic reads substantially increased gene detection to 7000 genes per nucleus. In addition, intronic reads were more frequently derived from long genes that are known to have brain-specific expression (Gabel et al. 2015) and that help define neuronal connectivity and signaling. Intronic reads may also reflect other cell-type specific features, such as retained introns or alternative isoforms. For example, intron retention provides a mechanism for the nuclear storage and rapid translation of long transcripts in response to neuronal activity (Mauger, Lemoine, and Scheiffele 2016).

We found that nuclei contain at least 20% of all cellular transcripts, and this percentage varies among cell types. Two small pyramidal neuron types have large nuclei relative to cell size that contain more than half of all transcripts. We detect 4000 more genes in single cells than single nuclei, but the majority of genes are detected equally well in both. Cytoplasm-enriched transcripts are missed by profiling single nuclei but include mostly house-keeping genes and pseudogenes, which are not related to neuronal identity. Nucleus-enriched transcripts include protein-coding and non-coding genes that are more likely to be cell-type markers than cytoplasmic transcripts. Overall, single cells do provide somewhat better detection of cell-type marker genes, thereby resulting in slightly better cluster separation for two pairs of highly similar cell types. Therefore, as more nuclei and cells are profiled, it is possible that finer discrimination of cell types may require single cell profiling. However, the benefits of profiling single nuclei may outweigh potential loss in the finest cell type resolution.

snRNA-seq is well suited for large-scale surveys of cellular diversity in various tissues and has the potential to be less cell-type biased. For example, single cell profiling of adult human cortex isolated more interneurons than excitatory neurons (Darmanis et al. 2015), whereas single nucleus profiling of the same tissue type isolated 30% interneurons and 70% excitatory neurons (Lake et al. 2016), close to the proportions found *in situ*. snRNA-seq also enables the use of stored frozen specimens to study cell types that will inform our understanding of human diversity and disease. As large scale initiatives begin to characterize transcriptomic cell types in the whole brain (Ecker et al. 2017) and whole organism (Regev et al. 2017), it is important to understand the strengths and limitations of each mRNA profiling technique.

## Materials and Methods

### Tissue preparation

Tissue samples were obtained from adult (postnatal day (P) 53-59)) male and female transgenic mice carrying a Cre transgene and a Cre-reporter transgene. Mice were anesthetized with 5% isoflurane and intracardially perfused with either 25 or 50 ml of ice cold, oxygenated artificial cerebral spinal fluid (ACSF) at a flow rate of 9 ml per minute until the liver appeared clear, or the full volume of perfusate had been flushed through the vasculature. The ACSF solution consisted of 0.5mM CaCl_2_, 25mM D-Glucose, 98mM HCl, 20mM HEPES, 10mM MgSO_4_, 1.25mM NaH_2_PO_4_, 3mM Myo-inositol, 12mM N-acetylcysteine, 96mM N-methyl-D-glucamine, 2.5mM KCl, 25mM NaHCO_3_, 5mM sodium L-Ascorbate, 3mM sodium pyruvate, 0.01mM Taurine, and 2mM Thiourea. The brain was then rapidly dissected and mounted for coronal slice preparation on the chuck of a Compresstome VF-300 vibrating microtome (Precisionary Instruments). Using a custom designed photodocumentation configuration (Mako G125B PoE camera with custom integrated software), a blockface image was acquired before each section was sliced at 250 µm intervals. The slice was then hemisected along the midline, and both hemispheres were then transferred to chilled, oxygenated ACSF.

Each slice-hemisphere was transferred into a Sylgard-coated dissection dish containing 3 ml of chilled, oxygenated ACSF. Brightfield and fluorescent images between 4X and 20X were obtained of the intact tissue with a Nikon Digital Sight DS-Fi1 or a Sentech STC-SC500POE camera mounted to a Nikon SMZ1500 dissecting microscope. To guide anatomical targeting for dissection, boundaries were identified by trained anatomists, comparing the blockface image and the slice image to a matched plane of the Allen Reference Atlas. In general, three to five slices were sufficient to capture the targeted region of interest, allowing for expression analysis along the anterior/posterior axis. The region of interest was then dissected and both brightfield and fluorescent images of the dissections were acquired for secondary verification. The dissected regions were transferred in ACSF to a microcentrifuge tube, and stored on ice. This process was repeated for all slices containing the target region of interest, with each region of interest deposited into a new microcentrifuge tube.

For whole cell dissociation, after all regions of interest were dissected, the ACSF was removed and 1 ml of a 2 mg/ml pronase in ACSF solution was added. Tissue was digested at room temperature (approximately 22°C) for a duration that consisted of adding 15 minutes to the age of the mouse (in days; *i*.*e*., P53 specimen had a digestion time of 68 minutes). After digestion, the pronase solution was removed and replaced by 1 ml of ACSF supplemented with 1% Fetal Bovine Serum (FBS). The tissue was washed two more times with the same solution and the sample was then triturated using fire-polished glass pipettes of decreasing bore sizes (600, 300, and 150 mm). The cell suspension was incubated on ice in preparation for fluorescence-activated cell sorting (FACS). FACS preparation involved adding 4’-6-diamidino-2-phenylindole (DAPI) at a final concentration of 4 µg/ml to label dead (DAPI+) versus live (DAPI-) cells. The suspension was then filtered through a fine-mesh cell strainer to remove cell aggregates. Cells were sorted by excluding DAPI positive events and debris, and gating to include red fluorescent events (tdTomato-positive cells). Single cells were collected into strip tubes containing 11.5µl of collection buffer (SMART-Seq v4 lysis buffer 0.83x, Clontech #634894), RNase Inhibitor (0.17U/ml), and ERCCs (External RNA Controls Consortium, MIX1 at a final dilution of 1x10-8) (Baker et al. 2005; Risso et al. 2014). After sorting, strip tubes containing single cells were centrifuged briefly and then stored at -80°C.

For nuclei isolation, dissected regions of interest were transferred to microcentrifuge tubes, snap frozen in a slurry of dry ice and ethanol, and stored at -80°C until the time of use. To isolate nuclei, frozen tissues were placed into a homogenization buffer that consisted of 10mM Tris pH 8.0, 250mM sucrose, 25mM KCl, 5mM MgCl2, 0.1% Triton-X 100, 0.5% RNasin Plus RNase inhibitor (Promega), 1X protease inhibitor (Promega), and 0.1mM DTT. Tissues were placed into a 1ml dounce homogenizer (Wheaton) and homogenized using 10 strokes of the loose dounce pestle followed by 10 strokes of the tight pestle to liberate nuclei. Homogenate was strained through a 30µm cell strainer (Miltenyi Biotech) and centrifuged at 900xg for 10 minutes to pellet nuclei. Nuclei were then resuspended in staining buffer containing 1X PBS supplemented with 0.8% nuclease-free BSA and 0.5% RNasin Plus RNase inhibitor. Mouse anti-NeuN antibody (EMD Millipore, MAB377, Clone A60) was added to the nuclei at a final dilution of 1:1000 and nuclei suspensions were incubated at 4°C for 30 minutes. Nuclei suspensions were then centrifuged at 400xg for 5 minutes and resuspended in clean staining buffer (1X PBS, 0.8% BSA, 0.5% RNasin Plus). Secondary antibody (goat anti-mouse IgG (H+L), Alexa Fluor 594 conjugated, ThermoFisher Scientific) was applied to nuclei suspensions at a dilution of 1:5000 for 30 minutes at 4°C. After incubation in secondary antibody, nuclei suspensions were centrifuged at 400xg for 5 minutes and resuspended in clean staining buffer. Prior to FACS, DAPI was applied to nuclei suspensions at a final concentration of 0.1µg/ml and nuclei suspensions were filtered through a 35µm nylon mesh to remove aggregates. Single nuclei were captured by gating on DAPI-positive events, excluding debris and doublets, and then gating on Alexa Fluor 594 (NeuN) signal. Strip tubes containing FACS isolated single nuclei were then briefly centrifuged and frozen at -80°C.

### RNA amplification and library preparation for RNA-seq

The SMART-Seq v4 Ultra Low Input RNA Kit for Sequencing (Clontech #634894) was used per the manufacturer’s instructions for reverse transcription of single cell RNA and subsequent cDNA synthesis. Single cells were stored in 8-strips at -80°C in 11.5 µl of collection buffer (SMART-Seq v4 lysis buffer at 0.83x, RNase Inhibitor at 0.17 U/µl, and ERCC MIX1 at a final dilution of 1x10-8 dilution). Twelve to 24 8-well strips were processed at a time (the equivalent of 1-2 96-well plates). At least 1 control strip was used per amplification set, containing 2 wells without cells but including ERCCs, 2 wells without cells or ERCCs, and either 4 wells of 10 pg of Mouse Whole Brain Total RNA (Zyagen, MR-201) or 2 wells of 10 pg of Mouse Whole Brain Total RNA (Zyagen, MR-201) and 2 wells of 10 pg Control RNA provided in the Clontech kit. Mouse whole cells were subjected to 18 PCR cycles after the reverse transcription step, whereas mouse nuclei were subjected to 21 PCR cycles. AMPure XP Bead (Agencourt AMPure beads XP PCR, Beckman Coulter A63881) purification was done using the Agilent Bravo NGS Option A instrument. A bead ratio of 1x was used (50 µl of AMPure XP beads to 50 µl cDNA PCR product with 1 µl of 10x lysis buffer added, as per Clontech instructions), and purified cDNA was eluted in 17 µl elution buffer provided by Clontech. All samples were quantitated using PicoGreen® on a Molecular Dynamics M2 SpectraMax instrument. A portion of the samples, and all controls, were either run on the Agilent Bioanalyzer 2100 using High Sensitivity DNA chips or the Advanced Analytics Fragment Analyzer (96) using the High Sensitivity NGS Fragment Analysis Kit (1bp-6000bp) to qualify cDNA size distribution. An average of 7.3 ng of cDNA was synthesized across all non-control samples. Purified cDNA was stored in 96-well plates at -20°C until library preparation.

Sequencing libraries were prepared using NexteraXT (Illumina, FC-131-1096) with NexteraXT Index Kit V2 Set A (FC-131-2001). NexteraXT libraries were prepared at 0.5x volume, but otherwise followed the manufacturer’s instructions. An aliquot of each amplified cDNA sample was first normalized to 30 pg/µl with Nuclease-Free Water (Ambion), then this normalized sample aliquot was used as input material into the NexteraXT DNA Library Prep (for a total of 75pg input). AMPure XP bead purification was done using the Agilent Bravo NGS Option A instrument. A bead ratio of 0.9x was used (22.5 ul of AMPure XP beads to 25 ul library product, as per Illumina protocol), and all samples were eluted in 22 µl of Resuspension Bu er (Illumina). All samples were run on either the Agilent Bioanalyzer 2100 using High Sensitivity DNA chips or the Advanced Analytics Fragment Analyzer (96) using the High Sensitivity NGS Fragment Analysis Kit (1bp-6000bp) to for sizing. All samples were quantitated using PicoGreen using a Molecular Dynamics M2 SpectraMax instrument. Molarity was calculated for each sample using average size as reported by Bioanalyzer or Fragment Analyzer and pg/µl concentration as determined by PicoGreen. Samples (5 µl aliquot) were normalized to 2-10 nM with Nuclease-free Water (Ambion), then 2 µl from each sample within one 96-index set was pooled to a total of 192 µl at 2-10 nM concentration. A portion of this library pool was sent to an outside vendor for sequencing on an Illumina HS2500. All of the library pools were run using Illumina High Output V4 chemistry. Covance Genomics Laboratory, a Seattle-based subsidiary of LabCorp Group of Holdings, performed the RNA-Sequencing services. An average of 229 M reads were obtained per pool, with an average of 2.0-3.1 M reads/cell across the entire data set.

### RNA-Seq data processing

Raw read (fastq) files were aligned to the GRCm38 mouse genome sequence (Genome Reference Consortium, 2011) with the RefSeq transcriptome version GRCm38.p3 (current as of 1/15/2016) and updated by removing duplicate Entrez gene entries from the gtf reference file for STAR processing. For alignment, Illumina sequencing adapters were clipped from the reads using the fastqMCF program (Aronesty 2011). After clipping, the paired-end reads were mapped using Spliced Transcripts Alignment to a Reference (STAR) (Dobin et al. 2013) using default settings. STAR uses and builds it own suffix array index which considerably accelerates the alignment step while improving sensitivity and specificity, due to its identification of alternative splice junctions. Reads that did not map to the genome were then aligned to synthetic constructs (i.e. ERCC) sequences and the *E.coli* genome (version ASM584v2). Quantification was performed using summer-izeOverlaps from the R package GenomicAlignments (Lawrence et al. 2013). Read alignments to the genome (exonic, intronic, and intergenic counts) were visualized as beeswarm plots using the R package *beeswarm*.

Expression levels were calculated as counts per million (CPM) of exonic plus intronic reads, and log_2_(CPM + 1) transformed values were used for a subset of analyses as described below. Gene detection was calculated as the number of genes expressed in each sample with CPM > 0. CPM values reflected absolute transcript number and gene length, i.e. short and abundant transcripts may have the same apparent expression level as long but rarer transcripts. Intron retention varied across genes so no reliable estimates of effective gene lengths were available for expression normalization. Instead, absolute expression levels were estimated as fragments per kilobase per million (FPKM) using only exonic reads so that annotated transcript lengths could be used.

### Selection of single nuclei and matched cells

463 of 487 (95%) of single nuclei isolated from layer 5 of mouse VISp passed quality control criteria: >500,000 genome-mapped reads, >75% reads aligned, and >50% unique reads. 12,866 single cells isolated from layers 1-6 of mouse VISp passed quality control criteria: >200,000 transcriptome mapped reads and >1000 genes detected (CPM > 0).

Gene expression was more likely to drop out in samples with lower quality cDNA libraries and for low expressing genes. To estimate gene dropouts due to stochastic transcription or technical artifacts (Kharchenko, Silberstein, and Scadden 2014), expression noise models were fit separately to single nuclei and cells using the “knn.error.models” function of the R package *scde* (version 2.2.0) with default settings and eight nearest neighbors. Noise models were used to calculate a dropout weight matrix that represented the likelihood of expression dropouts based on average gene expression levels of similar nuclei or cells using mode-relative weighting (“SCDE by Kharchenko Lab at Harvard DBMI”, n.d.). The probability of dropout for each sample (s) and gene (g) was estimated based on two expression measurements: average expected expression level of similar samples, 
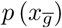
, and observed expression levels, *p* (*x_sg_*), using the “scde.failure.probability” and “scde.posteriors” functions. The dropout weighting was calculated as a combination of these probabilities: 
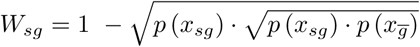
.

Dropout weighted Pearson correlations were calculated between all pairs of nuclei and cells using 42,003 genes expressed in at least one nucleus and one cell. The cell with the highest correlation to any nucleus was selected as the best match, and this cell and nucleus were removed from further analysis. This process was repeated until 463 best matching cells were selected, and the expression correlations were compared to correlations of the best matching pairs of nuclei (Figure 1B). The Cre-lines and dissected cortical layers of origin of the best matching cells were summarized as bar plots (Figure S1). Unweighted Pearson correlations were also calculated between all pairs of nuclei and cells to test the effect of accounting for dropouts on sample similarities (Figure 2B).

### Differential expression analysis

Gene detection was estimated as the proportion of cells and nuclei expressing each gene (CPM > 0). In order to estimate the expected variability of gene detection as a result of population sampling, cells were randomly split into two sets of 231 and 232 cells and genes were grouped into 50 bins based on detection in the first set of cells. For each bin of genes, the 97.5 percentile of detection was calculated for the second set of cells. A 95% confidence interval of gene detection was constructed by reflecting this these binned quantiles across the line of unity. Data were summarized with a hexagonal binned scatter plot and a log-transformed color scale using the R package *ggplot2 (Wickham 2009)*.

Differential expression between nuclei and cells was calculated with the R package limma (Ritchie et al. 2015) using default settings and log_2_(CPM + 1) expression defined based on two sets of reads: introns plus exons and only exons. Significantly differentially expressed were defined as having >1.5-fold change and a Benjamini-Hochberg corrected P-value < 0.05. Gene expression distributions of nuclei or cells within a cluster were visualized using violin plots, density plots rotated 90 degrees and reflected on the Y-axis.

Differences in alignment statistics and gene counts were calculated between cells, nuclei, and total RNA controls (or just cells and nuclei) with analysis of variance using the “aov” function in R (Chambers, Freeny, and Heiberger 1992). P-values for all comparisons were P<10^−13^.

Two sets of nucleus- and cell-enriched genes (introns plus exons and exons only) were tested for gene ontology (GO) enrichment using the ToppGene Suite (Chen et al. 2009). Significantly enriched (Benjamini-Hochberg false discovery rate < 0.05) GO terms were summarized as tree maps with box sizes proportional to -log10(P-values) using REVIGO (Supek et al. 2011) (Figure S2).

### Clustering

Nuclei and cells were grouped into transcriptomic cell types using an iterative clustering procedure based on community detection in a nearest neighbor graph as described in (Levine et al. 2015). Clustering was performed using gene expression quantified with exonic reads only or intronic plus exonic reads for two key clustering steps: selecting significantly variable genes and calculating pairwise similarities between nuclei. Four combinations of expression quantification for nuclei and cells resulted in eight independent clustering runs.

For each gene, log_2_(CPM + 1) expression was centered and scaled across samples. Noise models were used to select significantly variable genes (adjusted variance > 1.25). Dimensionality reduction was performed with principal components analysis (PCA) on variable genes, and the covariance matrix was adjusted to account for gene dropouts using the product of dropout weights across genes for each pair of samples. A maximum of 20 principal components (PCs) were retained for which more variance was explained than the broken stick null distribution, a conservative method of PC retention (Jackson 1993).

Nearest-neighbor distances between all samples were calculated using the “nn2” function of the R package *RANN*, and Jaccard similarity coefficients between nearest-neighbor sets were computed. Jaccard coefficients measured the proportion of nearest neighbors shared by each sample and were used as edge weights in constructing an undirected graph of samples. Louvain community detection was used to cluster this graph with 15 nearest neighbors. Considering more than 15 neighbors reduced the power to detect small clusters due to the resolution limit of community detection (Fortunato and Barthelemy 2007). Considering fewer than 15 neighbors increased over-splitting, as expected based on simulations by (Reichardt and Bornholdt 2006). Fewer nearest neighbors were used only when there were 15 or fewer samples total.

Clustering significance was tested by comparing the observed modularity to the expected modularity of an Erdös-Rényi random graph with a matching number of nodes and average connection probability. Expected modularity was calculated as the maximum estimated by two reported equations (Guimerà, Sales-Pardo, and Amaral 2004; Reichardt and Bornholdt 2006). Samples were split into clusters only if the observed modularity was greater than the expected modularity, and only clusters with distinct marker genes were retained. Marker genes were defined for all cluster pairs using two criteria: 1) significant differential expression (Benjamini-Hochberg false discovery rate < 0.05) using the R package *limma* and 2) either binary expression (CPM > 1 in >50% samples in one cluster and <10% in the second cluster) or >100-fold difference in expression. Pairs of clusters were merged if either cluster lacked at least one marker gene.

Clustering was applied iteratively to each sub-cluster until the occurrence of one of four stop criteria: 1) fewer than six samples (due to a minimum cluster size of three); 2) no significantly variable genes; 3) no significantly variable PCs; 4) no significant clusters.

To assess the robustness of clusters, the iterative clustering procedure described above was repeated 100 times for random sets of 80% of samples. A co-clustering matrix was generated that represented the proportion of clustering iterations that each pair of samples were assigned to the same cluster. Average-linkage hierarchical clustering was applied to this matrix followed by dynamic branch cutting using “cutreeHybrid” in the R package *WGCNA* (Langfelder, Zhang, and Horvath 2007) with cut height ranging from 0.01 to 0.99 in steps of 0.01. A cut height was selected that resulted in the median number of clusters detected across all 100 iterations. Cluster cohesion (average within cluster co-clustering) and separation (difference between within cluster co-clustering and maximum between cluster co-clustering) was calculated for all clusters. Marker genes were defined for all cluster pairs as described above, and clusters were merged if they had a co-clustering separation <0.25 or either cluster lacked at least one marker gene.

### Scoring marker genes based on cluster specificity

Many genes were expressed in the majority of nuclei or cells in a subset of clusters. A marker score (beta) was defined for all genes to measure how binary expression was among clusters, independent of the number of clusters labeled. First, the proportion (*x_i_*) of samples in each cluster that expressed a gene above background level (CPM > 1) was calculated. Then, scores were defined as the squared differences in proportions normalized by the sum of absolute differences plus a small constant (ε) to avoid division by zero. Scores ranged from 0 to 1, and a perfectly binary marker had a score equal to 1.

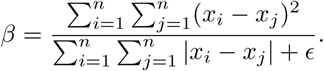

### Cluster dendrograms

Clusters were arranged by transcriptomic similarity based on hierarchical clustering. First, the average expression level of the top 1200 marker genes (i.e. highest beta scores) was calculated for each cluster. A correlation-based distance matrix (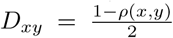) was calculated, and complete-linkage hierarchical clustering was performed using the “hclust” R function with default parameters. The resulting dendrogram branches were reordered to show inhibitory clusters followed by excitatory clusters, with larger clusters first, while retaining the tree structure. Note that this measure of cluster similarity is complementary to the co-clustering separation described above. For example, two clusters with similar gene expression patterns but a few binary marker genes may be close on the tree but highly distinct based on co-clustering.

### Matching clusters based on marker gene expression

Nuclei and cell clusters were independently compared to published mouse VISp cell types (Tasic et al. 2016). The proportion of nuclei or cells expressing each gene with CPM > 1 was calculated for all clusters. Approximately 400 genes were markers in both data sets (beta score > 0.3) and were expressed in the majority of samples of between one and five clusters. Markers expressed in more than five clusters were excluded to increase the specificity of cluster matching. Weighted correlations were calculated between all pairs of clusters across these genes and weighted by beta scores to increase the influence of more informative genes. Heatmaps were generated to visualize all cluster correlations. All nuclei and cell clusters had reciprocal best matching clusters from Tasic et al. and were labeled based on these reported cluster names.

Next, nuclei and cell clusters were directly compared using the above analysis. All 11 clusters had reciprocal best matches that were consistent with cluster labels assigned based on similarity to published types. The most highly conserved marker genes of matching clusters were identified by selecting genes expressed in a single cluster (>50% of samples with CPM > 1) and with the highest minimum beta score between nuclei and cell clusters. Two additional marker genes were identified that discriminated two closely related clusters. Violin plots of marker gene expression were constructed with each gene on an independent, linear scale.

Nuclei and cell clusters were also compared by calculating average cluster expression based only on intronic or exonic reads and calculating a correlation-based distance using the top 1200 marker genes as described above. Hierarchical clustering was applied to all clusters quantified using the two sets of reads. In addition, the average log2(CPM + 1) expression across all nuclei and cells was calculated using intronic or exonic reads.

Cluster separation was calculated for individual nuclei and cells as the average within cluster co-clustering of each sample minus the maximum average between cluster co-clustering. Separations for matched pairs of clusters were visualized with box plots and compared using a Student’s t-test, and significance was tested after Bonferroni correction for multiple testing. Finally, a linear model was fit to beta marker scores for genes that were expressed in at least one but not all cell and nuclear clusters, and the intercept was set to zero.

### Estimating proportions of nuclear transcripts

The nuclear proportion of transcripts was estimated in two ways. First, all intronic reads were assumed to be from transcripts localized to the nucleus so that the proportion of intronic reads measured in cells should decrease linearly with the nuclear proportion of the cell as nuclear reads are diluted with cytoplasmic reads. For each cell type, the nuclear proportion was estimated as the proportion of intronic reads in cells divided by the proportion of intronic reads in matched nuclei. Second, the nuclear proportion was estimated as the average ratio of cell to nuclear expression (CPM) using only exonic reads of three highly expressed nuclear genes (*Snhg11*, *Malat1*, and *Meg3*). The standard deviation of nuclear proportion estimates were calculated based on standard error propagation of variation in intronic read proportions and expression levels. Nuclear proportion estimates were compared with linear regression, and the estimate based on relative expression levels was used for further analysis.

The nuclear proportion of transcripts for all genes was estimated for each cell type as the ratio of average expression (CPM) in nuclei versus matched cells multiplied by the nuclear proportion of all transcripts. Estimated proportions greater than 1 were set equal to 1 for each cell type, and a weighted average proportion was calculated for each gene with weights equal to the average log_2_(CPM + 1) expression in each cell type. 11,932 genes were expressed in at least one nuclear or cell cluster (>50% samples expressed with CPM > 1) and were annotated as one of three gene types – protein-coding, protein non-coding, or pseudogene – using gene metadata from NCBI (ftp://ftp.ncbi.nlm.nih.gov/gene/DATA/GENE_INFO/Mammalia/Mus_musculus.gene_info.gz; downloaded 10/12/2017). For each type, histograms of gene counts with different nuclear proportions were generated. Next, beta marker score distributions were visualized as violin plots, and differences across gene types were compared with a Kruskal-Wallis rank sum test followed by Wilcoxon signed rank unpaired tests. Finally, genes were grouped into 10 bins of estimated nuclear proportions, from high cytoplasmic enrichment to high nuclear enrichment, and beta marker score distributions were visualized as box plots. A linear regression was fit to marker scores versus nuclear proportion.

Nuclear transcript proportions were compared to nuclear proportions estimated for mouse liver and pancreatic beta cells based on data from (Halpern et al. 2015). Ratios of normalized nuclear and cytoplasmic transcript counts were calculated in four tissue replicates. Average ratios were calculated for genes with at least one count in either fraction in at least one tissue. Nuclear proportion estimates for all genes with data from both data sets (n = 4373) were compared with Pearson correlation, a linear model with intercept set equal to zero, and histograms with a bin width of 0.02.

### Colorimetric *in situ* hybridization

*In situ* hybridization data for mouse cortex was from the Allen Mouse Brain Atlas (Lein et al. 2007). All data is publicly accessible through www.brain-map.org. Data was generated using a semiautomated technology platform as described in (Lein et al. 2007). Mouse ISH data shown is from primary visual cortex (VISp) in the Paxinos Atlas (Paxinos and others 2013).

### Multiplex fluorescence RNA *in situ* hybridization and quantification of nuclear versus cytoplasmic transcripts

The RNAscope multiplex fluorescent kit was used according to the manufacturer’s instructions for fresh frozen tissue sections (Advanced Cell Diagnostics), with the exception that 16µm tissue sections were fixed with 4% PFA at 4°C for 60 minutes and the protease treatment step was shortened to 15 minutes at room temperature. Probes used to identify nuclear and cytoplasmic enriched transcripts were designed antisense to the following mouse genes: *Calb1*, *Grik1*, and *Pvalb*. Following hybridization and amplification, stained sections were imaged using a 60X oil immersion lens on a Nikon TiE epifluorescence microscope.

To determine if spots fell within the nucleus or cytoplasm, a boundary was drawn around the nucleus to delineate its border using measurement tools within Nikon Elements software. To delineate the cytoplasmic boundary of each cell, a circle with a diameter of 15um was drawn and centered over the cell (Fig. 5). RNA spots in each channel were quantified manually using counting tools available in the Nikon Elements software. Spots that fell fully within the interior boundary of the nucleus were classified as nuclear transcripts. Spots that fell outside of the nucleus but within the circle that defined the cytoplasmic boundary were classified as cytoplasmic transcripts. Additionally, if spots intersected the exterior boundary of the nucleus they were classified as cytoplasmic transcripts. To prevent double counting of spots and ambiguities in assigning spots to particular cells, labeled cells whose boundaries intersected at any point along the circumference of the circle delineating their cytoplasmic boundary were excluded from the analysis. A linear regression was fit to nuclear versus soma probe counts, and the slope was used to estimate the nuclear proportion.

### *In situ* quantification of nucleus and soma size

Coronal brain slices from *Nr5a1-Cre;Ai14*, *Scnn1a-Tg3-Cre;Ai14*, and *Rbp4-Cre KL100;Ai14* mice were stained with anti-dsRed (Clontech #632496) to enhance tdTomato signal in red channel and DAPI to label nuclei. Maximum intensity projections from six confocal stacks of 1-mm intervals were processed for analysis. Initial segmentation was performed by CellProfiler (Lamprecht, Sabatini, and Carpenter 2007) to identify nuclei from the DAPI signal and soma from the tdTomato signal. Segmentation results were manually verified and any mis-segmented nuclei or somata were removed or re-segmented if appropriate. Area measurement of segmented nuclei and somata was performed in CellProfiler in Layer 4 from *Nr5a1-Cre;Ai14* and *Scnn1a-Tg3-Cre;Ai14* mice, and in Layer 5 from *Rbp4-Cre_KL100;Ai14* mice. A linear regression was fit to nuclear versus soma area to highlight the differences between Cre-lines.

For measurements of nucleus and soma size agnostic to Cre driver, we used 16 µm-tissue sections from P56 mouse brain. To label nuclei, DAPI was applied to the tissue sections at a final concentration of 1mg/ml. To label cell somata, tissue sections were stained with Neurotrace 500/525 fluorescent Nissl stain (ThermoFisher Scientific) at a dilution of 1:100 in 1X PBS for 5 minutes, followed by brief washing in 1X PBS. Sections were coverslipped with Fluoromount-G (Southern Biotech) and visualized on a Nikon TiE epiuorescence microscope using a 40x oil objective. Soma and nuclei area measurements were taken by tracing the boundaries of the Nissl-stained soma or DAPI-stained nucleus, respectively, using cell measurement tools available in the Nikon TiE microscope software. All cells with a complete nucleus clearly present within the section were measured, except that we excluded glial cells which had very small nuclei and scant cytoplasm. Measurements were taken within a 40x field of view across an entire cortical column encompassing layers 1-6, and the laminar position of each cell (measured as depth from the pial surface) was tracked along with the nucleus and soma area measurements for each cell.

For each cell in the experiments above, the nuclear proportion was estimated as the ratio of nucleus and soma area raised to the 3/2 power. This transformation was required to convert area to volume measurements and assumed that the 3-dimensional geometries of soma and nuclei were reflected by their cross-sectional profiles. This is true for approximately symmetrical shapes such as most nuclei and some somata, but will lead to under- or over-estimates of nuclear proportions for asymmetrical cells. Therefore, the estimated nuclear proportion of any individual cell may be inaccurate, but the average nuclear proportion for many cells should be relatively unbiased.

### Code availability

Data and code to reproduce all figures are publicly available from GitHub at https://github.com/AllenInstitute/NucCellTypes.

## Acknowledgements

The authors thank the Allen Institute for Brain Science founders, P. G. Allen and J. Allen, for their vision, encouragement, and support.

## Supplemental Figures

**Figure S1:**
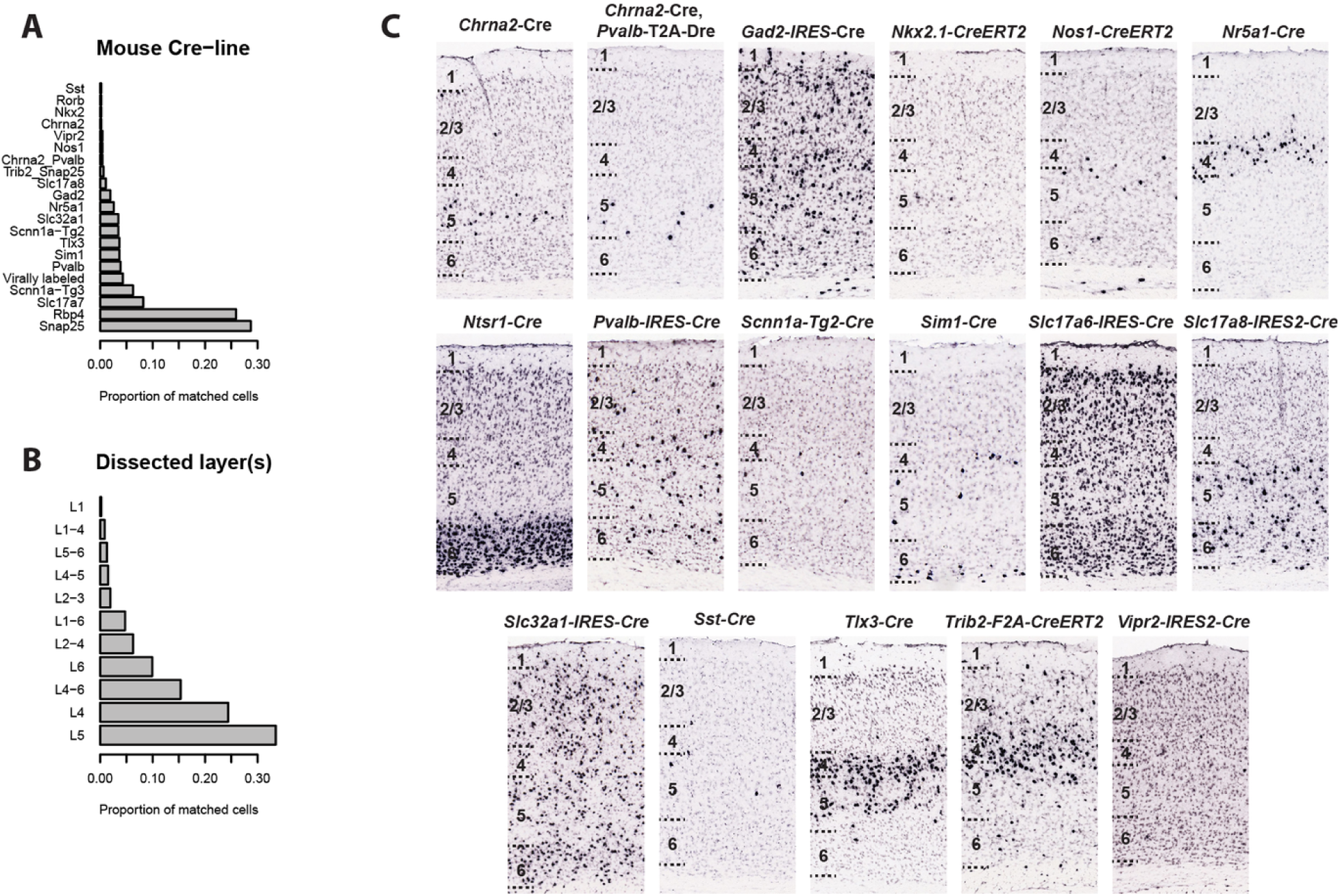
[Figure 1 - supplemental] Properties of 463 cells matched to nuclei. **(A)** Proportion of matched cells isolated from transgenic mouse lines that label different subsets of cortical neurons. Note that a small number of “virally labeled” cells (<5%) were FAC sorted from wild-type mice based on retrograde labeling by viral injections into various cortical and subcortical structures. **(B)** Proportion of matched cells dissected from one or more adjacent layers of cortex. **(C)** ISH images from additional mouse Cre-lines from which the best matching cells were most commonly derived. ISH images show all cortical layers within VISp.

**Figure S2:**
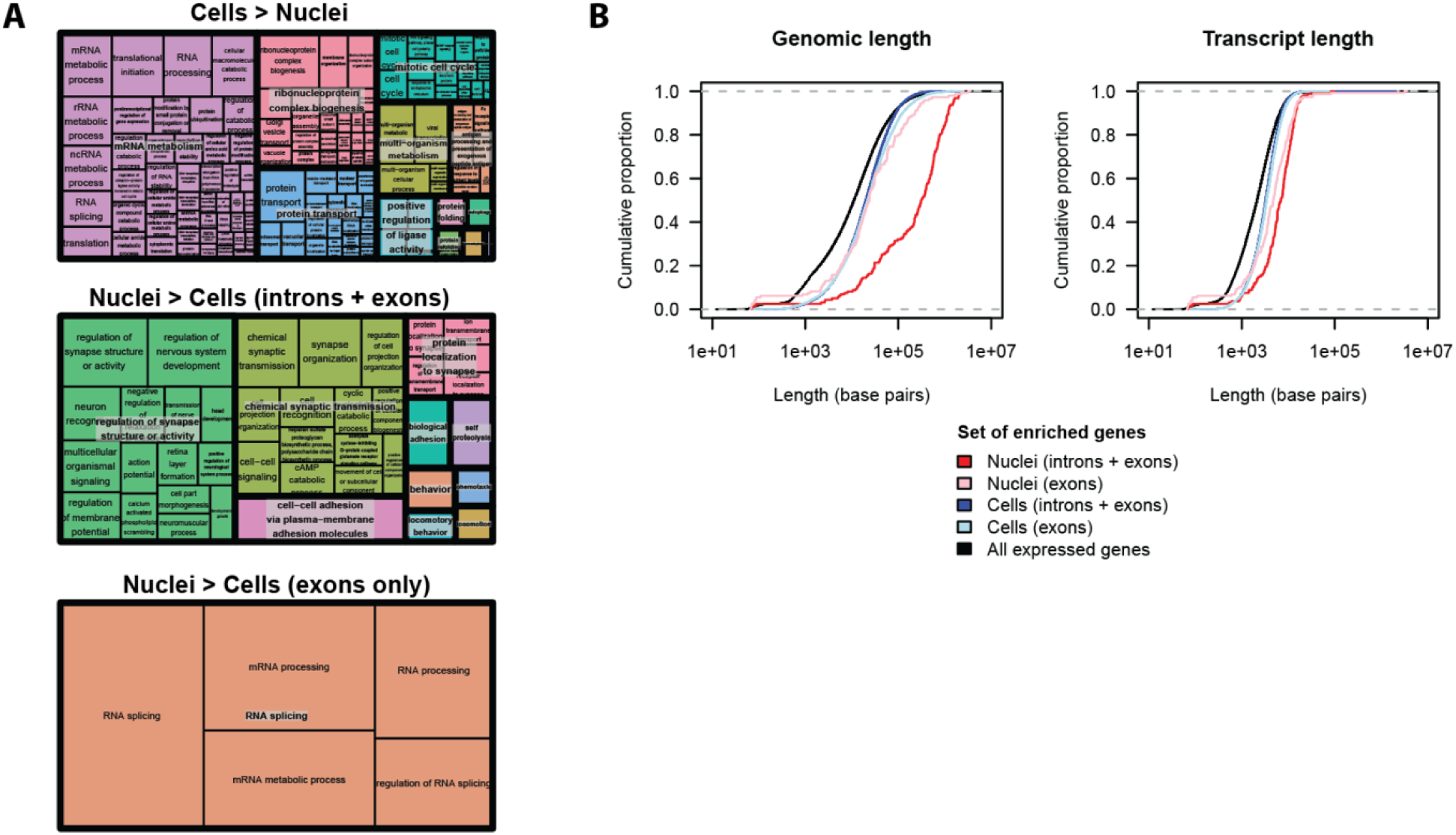
[Figure 2 - supplemental] Nuclear enrichment of transcripts related to neuron function can be explained by nuclear intron retention of long genes. **(A)** REVIGO (Supek et al. 2011) summaries of gene ontology (GO) enrichment of genes enriched in cells or nuclei. Including introns dramatically changes the functional categories of nuclear but not cell enriched genes. **(B)** Cumulative distribution of genomic and transcript lengths for genes enriched in nuclei and cells (fold change > 1.5) based on expression of exons or introns plus exons. Using introns plus exons, the median genomic length of nuclear enriched genes is 16-fold longer than cell enriched genes. Using exons only, there is no significant difference in genomic lengths (Kolmogorov-Smirnov test P-value = 0.27).

**Figure S3:**
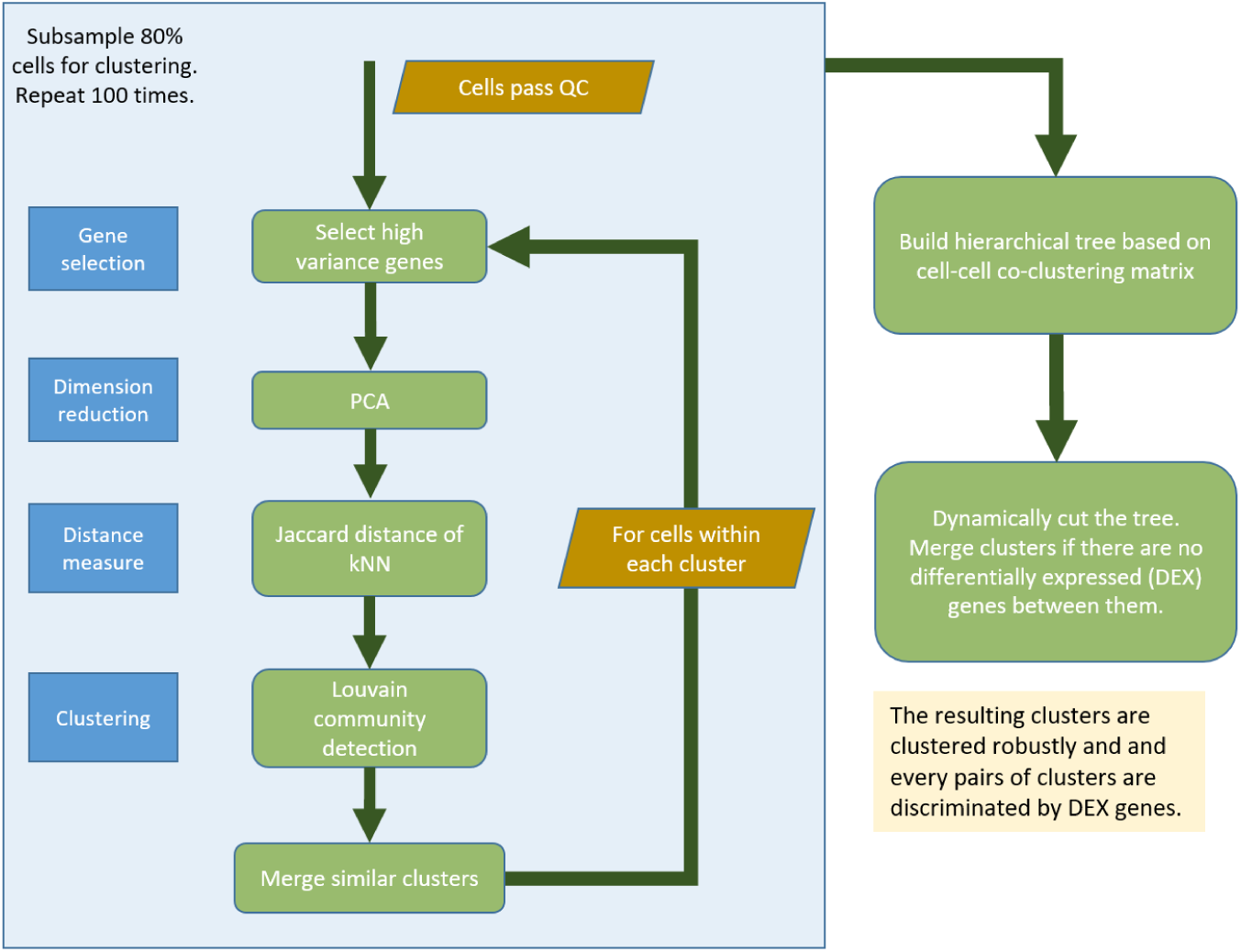
[Figure 3 - supplemental] Overview of single nucleus RNA-seq clustering pipeline. See methods for a detailed description of clustering steps.

**Figure S4:**
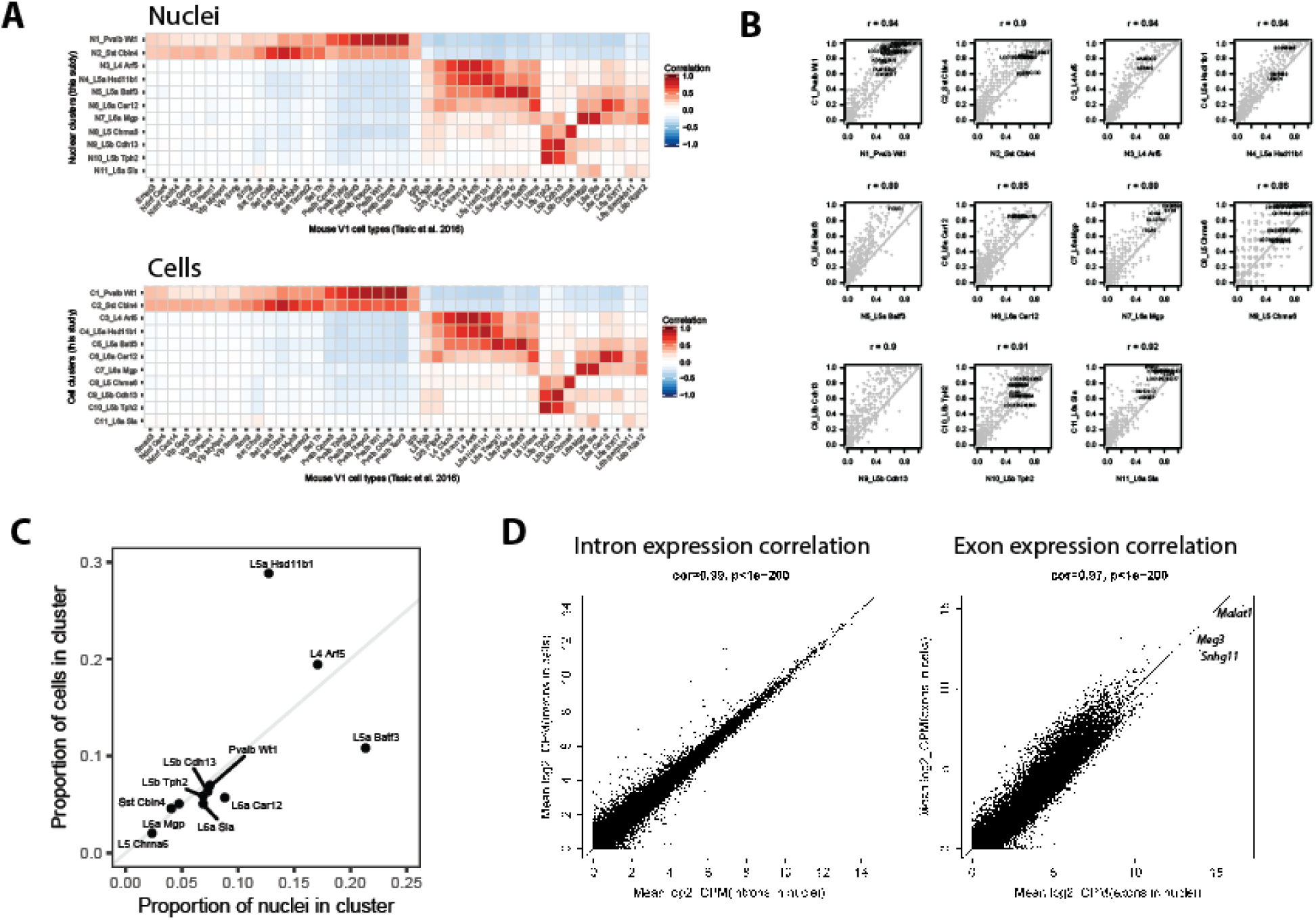
[Figure 4 - supplemental] Nuclear and cell clusters are well matched based on marker gene expression. **(A)** Pairwise correlations between previously reported mouse VISp cell type clusters (Tasic et al. 2016) and nuclear and cell clusters using average cluster expression of the top shared marker genes. Heatmaps show remarkably similar correlation patterns, supporting the existence of a well matched set of nuclear and cell clusters. Nuclear and cell clusters were annotated based on the reciprocal best matching published cluster name and mapped to two interneuron types and five of eight layer 5 excitatory neuron types. **(B)** Comparisons of the proportion of nuclei or cells expressing marker genes (CPM > 1) for matched pairs of clusters. Correlations are reported at the top of each scatter plot, and cell type specific markers are labeled. As expected based on Figure 2C, gene detection is consistently higher in cells than nuclei. **(C)** Matched clusters have similar proportions of nuclei and cells (except for two closely related cell types, L5a Hsd11b1 and L5 Batf3), which supports the accuracy of the initial correlation based mapping of single nuclei to cells. **(D)** Average gene expression quantified based on intronic reads is more highly correlated between cells and nuclei than expression quantified based on exonic reads, particularly for highly expressed genes. *Malat1*, *Meg3*, and *Snhg11* are the three highest expressing genes in nuclei and have consistently lower expression in cells, as expected based on their reported nuclear localization.

**Figure S5:**
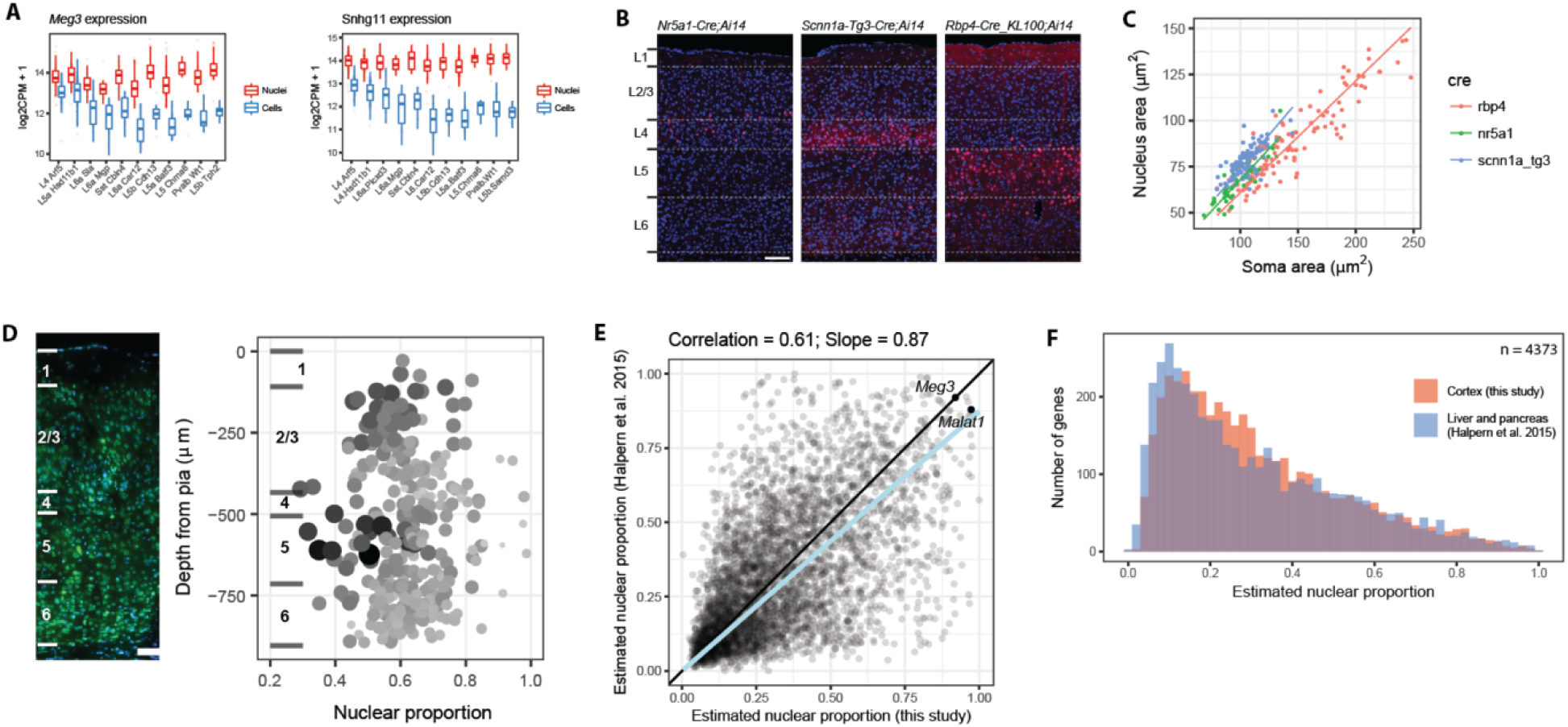
[Figure 5 - supplemental] Nuclear proportion estimates are supported by multiple genes and consistent with previously reported values. **(A)** Box plots of log_2_-transformed expression of two nuclear transcripts, *Meg3* and the small nucleolar RNA *Snhg11*, in matched nuclear and cell clusters. **(B)** Representative sections of VISp from three Cre-driver mouse lines with layer boundaries, nuclei labeled with DAPI (blue), and subsets of neurons labeled with tdTomato (red). Scale bar is 100 μm. **(C)** Nucleus and soma area measurements from three Cre-lines, and linear regressions to estimate nuclear proportions. **(D)** Left: Section of VISp from wild type mouse labeled with DAPI and Neurotrace 500 fluorescent Nissl stain with layer boundaries indicated by white lines. Scale bar is 100 μm. Right: Nuclear proportion was quantified based on nucleus and soma area measurements and plotted as a function of cortical depth. Size and darkness of points are proportional to soma area. **(E)** Average nuclear proportions of 4,373 genes (mostly house-keeping) also expressed in mouse pancreatic beta-cells and liver cells (Halpern et al. 2015) are moderately correlated with and approximately 13% less than estimated proportions in this study. **(F)** The distributions of nuclear proportions are highly similar with slightly higher reported cytoplasmic enrichment for reported genes. Note that the matched set of genes includes 99% protein-coding genes so the distributions more closely resemble those genes in Figure 5D.

## Supplemental Tables

**Table S1** [Figure 2 - supplemental]. Average gene expression and detection in matched nuclei and cells.

**Table S2** [Figure 2 - supplemental]. Differentially expressed genes in cells versus nuclei using intronic plus exonic reads.

**Table S3** [Figure 2 - supplemental]. Differentially expressed genes in cells versus nuclei using only exonic reads.

**Table S4** [Figure 2 - supplemental]. Gene ontology (GO) enrichment of differentially expressed genes in cells and nuclei.

**Table S5** [Figure 4 - supplemental]. Cre-driver line composition of cell clusters.

**Table S6** [Figure 5 - supplemental]. Gene properties including the number of clusters with any expression, maximum cluster expression, cell type marker score, and estimated nuclear proportion of transcripts.

